# Selection and neutral mutations drive pervasive mutability losses in long-lived B cell lineages

**DOI:** 10.1101/163741

**Authors:** Marcos Costa Vieira, Daniel Zinder, Sarah Cobey

## Abstract

High-affinity antibodies arise within weeks of infection from the evolution of B cell receptors under selection to improve antigen recognition. This rapid adaptation is enabled by the frequency and distribution of highly mutable “hotspotx” motifs in B cell receptor genes. High mutability in antigen binding regions (CDRs) creates variation in binding affinity, whereas low mutability in structurally important regions (FRs) may reduce the frequency of destabilizing mutations. During the response, the loss of mutational hotspots and changes in their distribution across CDRs and FRs are predicted to compromise the adaptability of B cell receptors, yet the contributions of different mechanisms to gains and losses of hotspots remain unclear. We reconstructed changes in anti-HIV B cell receptor sequences and show that mutability losses were about 60% more frequent than gains in both CDRs and FRs, with the higher relative mutability of CDRs maintained throughout the response. At least 34% of the mutability losses were caused by synonymous mutations. However, non-synonymous substitutions caused most of the mutability loss in CDRs. Because CDRs also show strong positive selection, this result suggests positive selection contributed to as much as 66% of the mutability loss in those regions. Although recurrent adaptation to the evolving virus could indirectly select for high mutation rates, we found no evidence of indirect selection to increase or retain hotspots. Our results suggest mutability losses are intrinsic to the neutral and adaptive evolution of B cell populations and might constrain their adaptation to rapidly evolving pathogens such as HIV and influenza.

## Introduction

High-affinity antibodies arise during the adaptive immune response from the very process that gave vertebrates an adaptive immune system in the first place: adaptation by natural selection. In response to infection or vaccination, mutagenic enzymes and error-prone polymerases cause somatic hypermutation of B cell receptors and thus create variation in their ability to bind antigen [1]. B cells with high-affinity receptors are more likely to receive survival and replication signals from helper T cells, and thus selection for improved antigen binding drives the development of high-affinity B cell receptors that are later secreted as antibodies [2–5]. Understanding how the immune system evolved to facilitate the rapid adaptation of B cell receptors during infection may provide general insights into the evolution of adaptability.

Two features of B cell receptor genes suggest their long-term evolution has been shaped by selection for adaptability during infection. First, the germline genes that recombine to produce B cell receptors are enriched for nucleotide motifs that are targeted with high frequency by the mutagenic enzymes involved in somatic hypermutation [6–8]. High mutation rates provide the genetic variation required for B cell adaptation, and low B cell mutation rates have been linked to immunodeficiency disorders [9, 10] and to the decline in immune function with age [11]. Second, mutational “hotspots” occur where mutations are most likely to be beneficial. Hotspots are concentrated in loops of the B cell receptor protein that are directly involved in antigen recognition (complementarity determining regions, CDRs) [12]. In contrast, structurally important regions of the B cell receptor (framework regions, FRs), which are usually less directly involved in antigen binding, are enriched with motifs that have low mutability [12]. In addition, mutations in CDRs are more likely to be non-synonymous and to produce non-conservative amino acid changes than mutations in FRs [13–16]. The differential mutability of CDRs and FRs appears to focus mutations to regions where they are likely to produce variation in antigen affinity without destabilizing the protein. The frequency and distribution of mutational hotspots in B cell receptor genes therefore seem to contribute to their adaptability during immune responses.

As B cells mutate during the immune response, however, changes in the frequency and distribution of mutational hotspots might affect the subsequent adaptability of B cell receptors. This change in adaptability may be especially important in B cell lineages that coevolve with pathogens like HIV and influenza. Experimental removal of hotspots decreases somatic hypermutation rates *in vitro* and in laboratory B cell lines [8]. Loss of highly mutable motifs has been hypothesized to occur during the immune response due to motifs’ propensity to mutate [17, 18], and decreased mutation rates due to such “hotspot decay” might explain declines in the evolutionary rates of B cell lineages over several years of HIV infection [19, 20]. In addition, changes in the distribution of highly mutable motifs across FRs and CDRs might increase the frequency of deleterious mutations in the former and decrease the frequency of beneficial mutations in the latter. However, evidence for consistent changes in mutability during the evolution of B cell lineages is inconclusive. Several metrics have been developed to estimate somatic hypermutation rates based on differences in the frequency of mutations across nucleotide motifs, and they range from the discrete classification of motifs into hotspots and non-hotspots [6, 7] to more recent models that quantify relative mutation rates on a continuous scale [12, 21, 22]. While the average mutability of B cell receptor sequences decreases during HIV infection when measured by the number of classically defined hotspots [17], long-term trends are less consistent for more recent metrics [20].

Factors other than the decay of hotspots through random mutations might affect the mutability of B cell receptors. First, highly mutable motifs might be regained through mutation. Second, selection for affinity and protein stability might favor non-synonymous mutations that incidentally increase or decrease mutability. Finally, selection might act on somatic hypermutation rates themselves. Although selection in theory can favor low mutation rates due to the reduced frequency of deleterious mutations [23, 24], rapidly changing environments may indirectly select for a higher mutation rate through its association with beneficial mutations [25–28]. Thus, although short-term differences in fitness among B cells arise from differences in the affinity of their receptors, B cells with more mutable CDRs might have a higher probability of producing high-affinity descendants able to keep up with evolving antigens in the long term. Indirect selection for mutability might therefore retain or increase the frequency of highly mutable motifs in CDRs, while mutability losses in FRs might be selected due to the reduced frequency of destabilizing mutations.

Understanding changes in mutability may reveal constraints on B cell adaptation, but the contributions of different mechanisms to changes in the frequency and distribution of mutational hotspots during B cell responses are largely unknown. We investigated the evolution of B cell receptor mutability by fitting phylogenetic models to sequences from long-lived anti-HIV B cell lineages. In characterizing mutability, we considered two sequence-based features that appear to have been strongly selected in the evolution of the adaptive immune system: overall mutability (the density of mutagenic nucleotide motifs) and also changes in the mutability of CDRs relative to FRs. First, we examined B cell mutability in the unmutated common ancestors of anti-HIV antibodies. Next, we investigated the effects of random mutations and positive selection for amino acid substitutions that increase affinity for antigen. Finally, we tested for selection to increase, retain or decrease the frequency of highly mutable motifs.

## Results

### Ancestral B cells have higher mutability in CDRs than FRs

To characterize changes in mutability during B cell evolution, we inferred the evolutionary histories of previously reported B cell lineages from three HIV-1 patients. The CH103 and VRC26 lineages comprise heavy and light chain B cell receptor sequences obtained from high-throughput sequencing over 144 and 206 weeks of infection in two patients, respectively [29, 30]. We also analyzed heavy chain sequences of three lineages from the VRC01 dataset, which was sampled from a third patient over a 15-year period [19]. The lineages we analyzed were originally investigated for having evolved the ability to neutralize diverse HIV strains.

To infer changes in mutability over time, we used Bayesian phylogenetic analyses (Methods) to obtain a sample of time-resolved trees from the posterior distribution of each lineage’s genealogy for the heavy and light chains separately, and to estimate the nucleotide sequences of all internal nodes. Mutabilities of the observed and the inferred internal sequences were estimated using the S5F model. This model assigns relative mutation rates to all five-nucleotide DNA motifs and is based on a large independent dataset of antigen-experienced B cells [21]. The mutability of each sequence was defined as the average S5F score across all sites in the B cell receptor sequence. We estimated the number, magnitude and distribution of mutability changes on all branches by contrasting the mutability of all pairs of parent-descendant nodes. Figure 1 illustrates mutability evolution in the heavy chain of lineage CH103.

**Fig 1.**
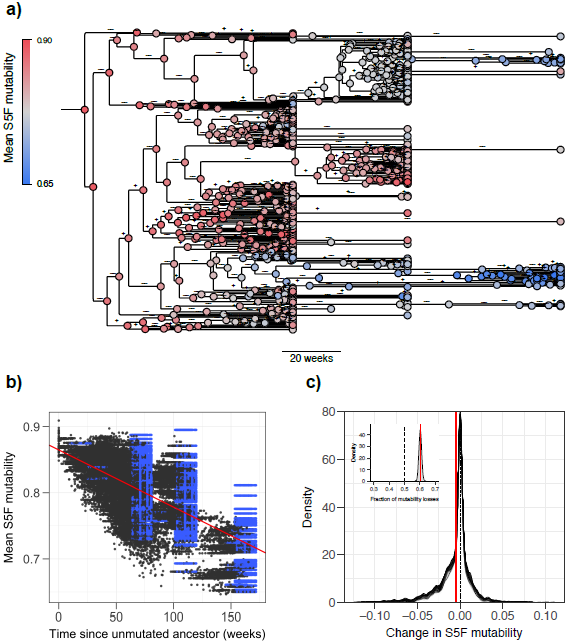
Evolution of S5F-mutability in the heavy-chain CH103 B cell lineage. a) Long-term declines in average mutability across the maximum clade-credibility tree. Nodes are colored according to the average S5F-mutability of the whole sequence. Branches measured in weeks are annotated to indicate gains (+) or losses (–) of mutability. b) Average mutability over time for a combined sample of 100 trees from the posterior distribution. Blue points correspond to terminal nodes (observed sequences), and black points correspond to inferred internal nodes. The red line represents an average of regression lines calculated for each tree in a sample of 1000 trees. c) Distribution of changes in mutability for a sample of 100 trees from the posterior distribution shown as overlaid densities. The inset plot shows the posterior distribution of the fraction of changes on a tree that were losses constructed from a sample of 1000 trees, with the 95% highest-posterior density interval shown in gray. Red lines indicate the means of the distributions.

We first characterized the mutability of each lineage’s unmutated ancestor. In the ancestors of all heavy and light chains, sites in CDRs had higher average mutability than sites in FRs. On average across the seven heavy and light chain ancestors, mutability was approximately 27% higher in CDRs than in FRs (range: 14-34%; Figure S1). Previous analyses of germline V genes (which, together with D and J genes, recombine to produce mature B cell receptors) showed that mutability was lower in FRs and higher in CDRs than expected based on their amino acid sequences, suggesting selection to increase the frequency of highly mutable motifs in CDRs and decrease their frequency in FRs [12, 15]. Consistent with those analyses, we found that the FRs of lineage ancestors had lower mutability than expected based on their amino acid sequences. On average across all heavy and light chains, the mean FR mutability of ancestral B cell receptors was lower than 99% of sequences obtained by randomizing their codons (according to usage frequencies in humans [31]) while keeping the amino acid sequences constant (range: 95-100%; Figure S1). However, while V genes often have higher CDR mutability than expected based on their amino acid sequences [12], mean CDR mutability in ancestral B cell receptors was, on average, greater than only 54% of randomized sequences with the same amino acid sequence (range: 18-95%, Figure S1). These patterns suggest that adaptation in these cells may on average be constrained more by destabilizing mutations in the FRs than a shortage of beneficial mutations in the CDRs.

### Mutability is more often lost than gained

Next we investigated long-term changes in mutability as B cell receptors evolved from their ancestors. Consistent with a previous analysis [17], average mutability decreased with time (Figure 1a,b, Figure S2). To investigate changes in mutability in greater detail, we analyzed the frequency of mutability gains and losses across branches. Mutability losses should arise from hotspot decay, positive selection of the amino acid changes that incidentally decrease mutability, selection for lower mutability due to the reduction in the frequency of deleterious mutations, or a combination of those factors. Mutability gains should reflect gains through mutation, positive selection of amino acid changes that incidentally increase mutability, indirect selection for higher mutation rates by association with beneficial mutations, or a combination of those factors. To summarize the net contributions of mutability-decreasing and mutability-increasing mechanisms, we computed the fraction of branches with mutability losses out of all branches with mutability changes. By computing the frequency of mutability losses for each tree in the posterior sample, we estimated its posterior distribution (Figure 1c).

Across the entire BCR sequence, mutability losses occurred more frequently than gains in four of the five lineages (Figure 1a,c, Figure 2). On average across the seven heavy and light chains, approximately 62% of changes in mutability were losses (range: 45% – 71%). Mutability losses were more frequent in the light chains (70% and 63%) than in the heavy chains (60% and 62%) of lineages CH103 and VRC26. The frequency of mutability was not significantly different from the frequency of gains in the heavy chain of VRC26 (95% highest posterior interval 48-74%) and was slightly lower than the frequency of gains in lineage VRC01-01 (42-49%).

**Fig 2.**
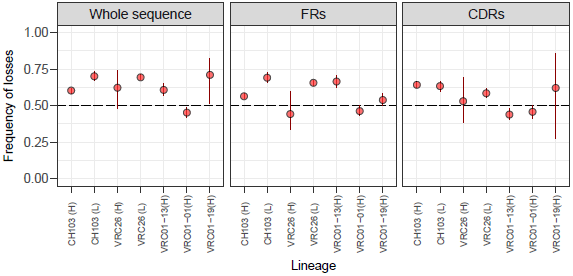
Frequency of losses relative to the total number of changes in S5F-mutability during the evolution of anti-HIV B cell lineages. Results are shown for the entire analyzed region of the B cell receptor, and separately for framework regions (FRs) and complementarity determining regions (CDRs). Each point denotes the fraction of changes in mutability that were losses, averaged across a sample of 1000 trees from the posterior distribution. Vertical red lines indicate the 95% highest-posterior density interval.

Long-term declines in mutability occurred both in FRs (five of seven heavy and light chains; Figure S3) and CDRs (six of seven heavy and light chains; Figure S4). Consistent with long-term declines in the average mutability of both regions, we found no long-term changes in the difference between CDR and FR mutabilities (Figure S5). Despite considerable amino acid divergence from the unmutated ancestors in FRs (16-67% for different heavy and light chains), sequences from the last sampling time in each dataset had lower FR mutability than 95% of randomizations with constant amino acid sequence (range: 70-99.9%, fig. S6). The lower mutability of ancestral FRs relative to their amino acid sequences was therefore retained throughout the evolution of the B cell lineages. CDRs, however, became less mutable relative to their amino acid sequences than the ancestors. On average across the seven heavy and light chains, CDRs of sequences from the last sampling times were more mutable than 28% of their corresponding randomizations (range: 8-64%, Figure S6), down from 54% in ancestral CDRs. Consistent with the net long-term trends, both CDRs and FRs had similar frequencies of mutability losses (FR average 57%, range: 44% – 69%; CDR average 56%, range: 44– 64%; Figure 2).

### Hotspot decay and selection for amino acid substitutions contribute to mutability losses

Highly mutable motifs have been hypothesized to decay due to their propensity to mutate [17, 18], but motifs should also be influenced by positive selection. The first case is straightforward. For instance, the hypothetical sequence CAG<u>C</u>TT contains the highly mutable cytosine at the center of the AGCTT motif [7, 8, 21], and a C→ T mutation in the underlined position would decrease the mutability of the site approximately 5-fold [21]. Positive selection on amino acid substitutions should influence motifs in two ways. Mutational hotspots can be disrupted if selection favors amino acid sequences whose codons happen to be, on average, less mutable than codons in the ancestral sequence. Selection can also affect mutability in neighboring codons. For example, if selection led to the replacement of CAG (glutamine) for CGG (arginine) in the sequence CAG<u>C</u>TT, the mutability of the underlined C nucleotide (not involved in the substitution) would decrease approximately 13-fold [21].

To evaluate the strength of positive selection in the B cell lineages, we used BASELINe [32] to quantify deviations in the frequency of non-synonymous substitutions from its expected value in the absence of selection (and under the mutational biases captured by the S5F model). In line with previous analyses of the same datasets [20] and of B cell lineages from healthy donors [33], we detected positive selection in the CDRs of four of the seven heavy and light chains (lineages CH103 and VRC26) and purifying selection in the FRs of all heavy and light chains. Framework regions had fewer non-synonymous substitutions than expected under neutral evolution, while CDRs had more non-synonymous substitutions than the neutral expectation (Figure 3). This result contrasts with repertoire-level analyses showing predominantly purifying selection across both types of regions [34].

**Fig 3.**
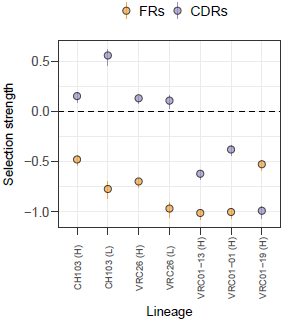
Selection in framework regions (FRs) and complementarity determining regions (CDRs) of B cell receptors from anti-HIV B cell lineages. Selection strength is measured as the log odds ratio between the observed ratio of non-synonymous to synonymous substitution frequencies, *π/*(1-*π*), and the ratio expected under the S5F mutability model in the absence of selection, 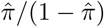 [21, 32]. Selection strength values greater than zero indicate positive selection, and values smaller than zero indicate purifying selection.

To estimate how much of the mutability losses in the CDRs is attributable to selection for amino acid substitutions, we partitioned inferred changes in mutability into changes caused by synonymous substitutions and changes caused by non-synonymous substitutions. Summed across all branches of the maximum-clade-credibility (MCC) trees, non-synonymous substitutions caused a net loss in the average CDR mutability of the same four heavy and light chains where positive selection occurred (Figure 4a). Similarly, synonymous substitutions caused a net decrease in the average CDR mutability of the four lineages whose CDRs were under positive selection. On average across those four lineages, non-synonymous substitutions accounted for approximately 66% of the inferred mutability loss in CDRs (range: 40% – 93%). On average, selection for amino acid substitutions may therefore have contributed to as much as 66% of the loss of mutability in the CDRs.

**Fig 4.**
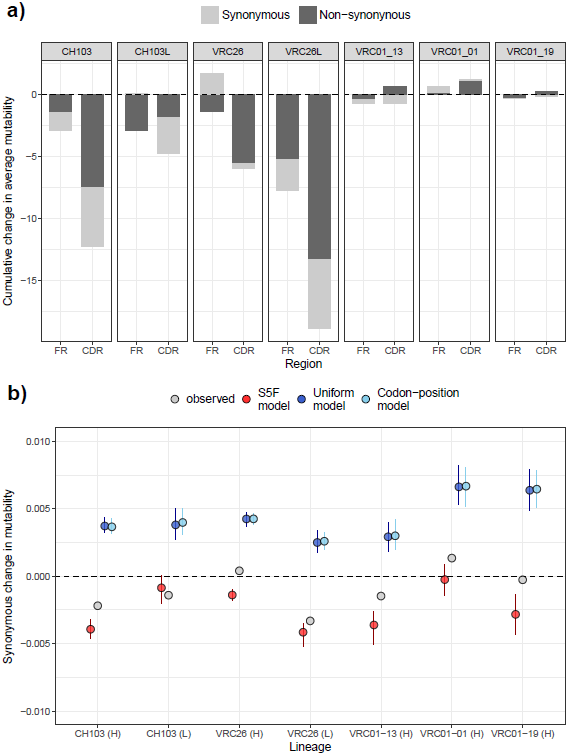
Changes in S5F mutability due to synonymous and non-synonymous substitutions in anti-HIV B cell lineages. A) Changes in mutability caused by synonymous and non-synonymous substitutions, summed across all branches of B cell genealogies. B) Changes in mutability due to synonymous substitutions, averaged across all branches. Gray indicates values inferred from the data, red indicates an S5F-based model where different nucleotide motifs mutate with different rates, dark blue indicates a model with no mutation rate variation across sites, and light blue indicates a model with different mutation rates for each position of a codon. Simulations were performed independently for each branch on the MCC tree of different anti-HIV B cell lineages, starting from the inferred sequence of the parent node. Each simulated sequence was constrained to have the same amino acid sequence as the inferred sequence for the descendant node. Vertical bars indicate the 95% range obtained from 100 simulations per model.

To test if the observed synonymous mutability losses were consistent with the decay of mutational hotspots under their predicted S5F mutability, we simulated B cell receptor evolution under a model that allows for variation in mutation rates across motifs based on their S5F mutability scores [21] and thus captures the propensity for certain motifs to mutate (Methods). We compared changes in mutability caused by synonymous substitutions simulated under the S5F-based model to changes under models that do not allow for motif-driven mutation rate variation. Instead, these models assume that the mutation rate is identical across all sites (“uniform” model) or depends on the position within a codon (“codon-position” model), with the relative rate for each position estimated from the data (Methods). For each branch on the MCC tree of each lineage, we started from the inferred nucleotide sequence of the branch’s parent node and simulated 100 replicates of a descendant sequence while constraining the simulations to produce the exact same amino acid substitutions inferred from the data. We also constrained simulated descendant sequences to have the same number of synonymous substitutions as inferred from the data but allowed them to occur at different positions.

Inferred mutability losses due to synonymous substitutions were consistent with the decay of mutational hotspots. On average across all branches, synonymous substitutions simulated under hotspot decay resulted in mutability losses in all seven heavy and light chains (Figure 4b). In contrast, synonymous substitutions simulated with constant mutation rates across motifs (uniform and codon-position models) increased mutability in all lineages.

### No evidence of indirect selection to retain mutability

Under persistent or recurrent selective pressures, alleles that increase the mutation rate may be indirectly selected due to their association with beneficial mutations [25–28]. Indirect selection for increased mutation rates might therefore lead to the conservation of highly mutable motifs in CDRs, where mutations that improve affinity are selected for during B cell evolution. In contrast, selection to reduce the frequency of deleterious mutations [23, 24] might directly favor mutability losses in FRs.

To test if mutability is subject to direct or indirect selection over the long term, we compared the frequency of synonymous changes in mutability between terminal and internal branches of the B cell trees. Terminal branches are expected to be enriched for deleterious mutations [35–38]. If mutability losses are deleterious in the long term, branches where mutability losses occurred should be less likely to contribute descendants to the B cell population, and would therefore be more likely terminal, rather than internal branches. In contrast, mutability losses should be more frequent on internal branches if mutability losses are beneficial. Because changes in mutability due to non-synonymous substitutions may be driven by selection for B cell receptor affinity and stability, we first restricted the analysis to mutability changes from synonymous substitutions.

We found no consistent evidence of selection to increase, retain, or decrease mutability (Figure 5). In one instance (the CDRs of light chain VRC26), mutability losses were more frequent on terminal branches than on internal branches, possibly indicating selection against mutability losses. In another instance (FRs of heavy chains VRC26), mutability losses were more frequent on internal branches than on internal branches, suggesting selection for mutability losses. In the remaining cases, the frequencies of synonymous mutability losses were similar in terminal and internal branches for both FRs and CDRs. The same lack of general trends was seen when we compared terminal branches, internal branches that belong to the main “backbone” of the tree, and other internal branches [38] (Figure S7), and also when we included mutability changes due to non-synonymous substitutions (results not shown).

**Fig 5.**
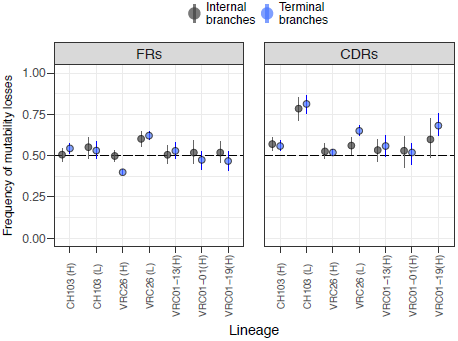
Frequency of losses relative to the total number of changes in whole-sequence S5F-mutability caused by synonymous substitutions during the evolution of anti-HIV B cell lineages. Blue indicates changes on terminal branches, and black indicates changes on internal branches. Results are shown separately for framework regions (FRs) and complementarity determining regions (CDRs). Each point denotes the frequency of changes in mutability that were losses, averaged across a sample of 1000 trees from the posterior distribution. Vertical lines indicate the 95% highest-posterior density interval.

### Results are consistent for three of four mutability metrics

In addition to the S5F model [21], we repeated the analyses using three other mutability metrics. The different mutability metrics were estimated from databases of somatic mutations not subject to the effect of selection, such as synonymous mutations, mutations in non-coding flanking regions of V genes, and mutations in unproductive B cell receptor genes that are not expressed by B cells but still undergo somatic hypermutation. Two of the alternative metrics are based on a discrete classification of DNA motifs into either hotspots or regular motifs: the number of WRCH and DGYW hotspots [7] or the number of “overlapping” (WGCW) hotspots [8], where *W* = {*A/T*}, *R* = {*A/G*}, *H* = {*A/T/C*}, *D* = {*A/T/G*} and *Y* = {*C/T*}. We also quantified mutability using the 7-mer model, which, similar to the S5F model, assigns relative mutation rates to different motifs on a continuous scale, but does so for seven-instead of five-nucleotide motifs [22].

Consistent with the results for S5F mutability, hotspots and overlapping hotspots decreased in number over time (Figure S8 and Figure S9) and were lost more frequently than gained (Figure S11). In contrast, average 7-mer mutability increased over time in four of five lineages (Figure S10), and losses of 7-mer mutability were approximately as frequent as gains across the entire B cell receptor sequence (fig. S11).

## Discussion

We used phylogenetic methods to reconstruct changes in B cell receptor sequences and found consistent loss of mutational hotspots over the course of the adaptive immune response. Our analyses shed light on previous studies of long-term trends in mutability [17, 20] by quantifying the contributions of different mechanisms to these losses. Selection for amino acid substitutions appears to have driven most of the mutability losses in the CDRs—precisely the regions where high mutation rates contribute the most to adaptation. However, mutability changes caused by synonymous substitutions throughout the sequence demonstrate mutability losses also arose from the spontaneous decay of highly mutable motifs through neutral mutations. We also show that both FRs and CDRs lost mutability frequently, and the higher mutability of CDRs relative to FRs observed in ancestral receptors persisted over at least several years of B cell evolution. Disruption of highly mutable motifs through selection and hotspot decay therefore seems to be intrinsic to the evolution of B cells during affinity maturation. These factors counteract the high mutation rate selected in the evolution of immunoglobulin genes, and they suggest adaptability may inevitably become compromised in evolving B cell lineages. However, the relatively high mutation rate in the CDRs compared to the FRs was preserved in these lineages, suggesting that some degree of adaptability may still be maintained. (These observations may reflect survival bias: lineages in which mutability differences between CDRs and FRs decrease over time may be less likely to persist in the long term.)

Theoretical simulations [26] and experimental evolution of bacteria [27, 28] suggest recurrent adaptation in asexual populations can select for increased mutation rates over the long term, but we found no consistent evidence of selection to increase or retain mutability during B cell evolution. Any long-term benefits of high mutation rates (in terms of increased chance of producing beneficial mutations) appear insufficient to overcome the rapid decay of highly mutable motifs through mutation and/or positive selection on amino acid substitutions. We also did not detect selection for reduced mutability in FRs, despite the potential for mutability losses to decrease the rate of destabilizing mutations. Because mutability is already low in the FRs of V genes [12, 15] and ancestral B cell receptors (this study), it is possible that the fitness effect of mutability losses in FRs is so small that they are nearly neutral given the effective size of the B cell population. Under this “drift-barrier” hypothesis [24], selection would therefore not be able to fix mutability losses.

Losses of mutational hotspots in CDRs may reduce the adaptability of antibody responses since lower mutation rates reduce the supply of genetic variation available for selection. Declines in mutation rates caused by hotspot losses may have contributed to reported declines in the evolutionary rates of broadly neutralizing B cell lineages during chronic HIV infection [20]. However, these losses may not be important for lineages that bind to conserved sites. As B cell receptors evolve high affinity for conserved sites, beneficial substitutions become rare, and the corresponding transition from positive to purifying selection should cause substitution rates to fall [20]. In line with the previous analysis [20], we found the longest-lived lineages to be under the strongest purifying selection in the CDRs. Those lineages also had minimal cumulative changes in mutability, suggesting most of the mutability loss occurred during early adaptation. However, as a result of hotspot loss, these lineages might adapt poorly if formerly conserved antigenic sites suddenly acquire mutations.

A direct link between motif-based mutability scores and absolute rates of B cell mutation in vivo is still lacking. Hotspot loss is associated with reduced substitution rates [19, 20], which are influenced not only by the underlying mutation rate but also by changes in selective pressures and generation times [20]. Using robust counting [39] and a random local clock model [40], we found no evidence for consistent declines in the synonymous or non-synonymous substitution rates of these lineages (Figures S12 to S15). Simulation-based power analysis (Supplementary Methods) revealed that synonymous and non-synonymous substitution rates had to decline at least 50% to be detected with this method (Figures S16 and S17). Observed long-term declines in average S5F mutability were less than 50% (Figure S2) and suggest their effects on substitution rates would not have been detected, assuming a one-to-one relationship between S5F mutability and the actual mutation rate. Additionally, inference of B cell genealogies may be affected by non-equilibrium nucleotide frequencies, distorting inferences of absolute rates [17]. Analyzing B cell sequences from controlled experimental infections with customized substitution models (e.g., [17, 34, 41]) could help quantify the relationship between mutability metrics and absolute mutation rates.

Our results suggest a trade-off between the short-term and long-term adaptability of antibody responses. Repeated infections by antigenically related pathogens, such as influenza viruses, often recall memory B cell populations [42–45]. Preferential recruitment of less mutable B cells at the expense of naive (or perhaps younger memory) B cells may compromise the long-term adaptability of the immune repertoire to these pathogens. Protection against such pathogens may rely on a robust naive response that can complement preexisting B cell lineages once they fail to adapt to new antigens. Finally, some strategies for eliciting universal responses against HIV and influenza attempt to recapitulate the evolution of long-lived, highly mutated B cell lineages that were found to produce broadly neutralizing antibodies in infected patients [46]. Our results suggest those approaches may be hindered by a decrease in the adaptability of highly mutated antibodies.

## Materials and Methods

### Sequence data

We analyzed BCR sequences from three published studies of B cell lineages sampled longitudinally in individual HIV-1 patients not subject to antiretroviral therapy. The CH103 lineage comprises broadly neutralizing antibodies (bnAbs) isolated from individual B cells and clonally related sequences bioinformatically isolated from high-throughput sequencing over 144 weeks of HIV-1 infection in a single donor [29]. Likewise, the VRC26 lineage comprises both bnAbs isolated from individually sorted B cells and clonally related heavy-chain sequences obtained from high-throughput sequencing over 206 weeks of HIV-1 infection in a different patient [30]. In both cases, BCR sequences obtained from high-throughput sequencing were classified as clonally related to the isolated antibodies based on V and J gene usage. A third lineage, VRC01, was sampled longitudinally from a third HIV patient for 15 years starting from approximately five years after the date of infection. VRC01 is a large lineage that possibly consists of multiple independent B cell lineages [19, 20]. We therefore used PARTIS, a recently developed hidden-markov-model-based method [47] to partition the heavy-chain VRC01 dataset into sets of sequences likely to have descended from the same naive B cell, and to identify their most likely germline V and J genes. We analyzed the three largest lineages identified in this way, hereafter VRC01-13, VRC01-01 and VRC01-19. We aligned each set of sequences in MACSE v1.01b [48] along with their concatenated V and J genes (for the heavy chains) or along with their V genes only (light chains) (Table 1). Including the J genes in the light chain datasets produced bad alignments across the J region.

**Table 1.** BCR alignments analyzed. The number of sequences includes the V+J or V germline sequences.

| Lineage (chain) | V gene | J gene | N sequences | N sites |
| --- | --- | --- | --- | --- |
| CH103 (heavy) | IGHV4-59*01 | IGHJ4*01 | 460 | 321 |
| CH103 (light) | IGLV3-1*01 | — | 175 | 285 |
| VRC26 (heavy) | IGHV3-30*18 | IGHJ3*01 | 681 | 489 |
| VRC26 (light) | IGLV1-51*02 | — | 464 | 294 |
| VRC01-13 (heavy) | IGHV1-2*04 | IGHJ1*01 | 157 | 351 |
| VRC01-01 (heavy) | IGHV-ORF15-1*04 | IGHJ4*03 | 124 | 372 |
| VRC01-19 (heavy) | IGHV1-2*04 | IGHJ1*01 | 110 | 438 |

### Phylogenetic inference

Using BEAST v.1.8.2 [49], we fit Bayesian phylogenetic models to the BCR sequence data in order to estimate time-resolved genealogies and internal node sequences for heavy and light chains separately. We used a GTR nucleotide substitution model, assumed a random local clock model to account for potential variation in substitution rates [40], and enabled robust counting [39] to estimate the numbers of synonymous and non-synonymous changes on each branch. To reduce the number of parameters, we used empirical base frequencies and assumed a shared nucleotide transition matrix across codon positions for the robust counting inference. The inferred dynamics of mutability losses and gains were qualitatively robust to the choice of demographic model (constant population size, logistic or exponential growth) used to calculate coalescent prior probabilities (Supplementary Methods). We set the V+J or V germline sequence of each alignment as the outgroup and assigned them a sampling time of zero to represent the assumption that, at the start of the infection, the ancestral BCR sequence was close to its corresponding germline genes (except for insertions and deletions at the junctions). For each dataset, four independent MCMC chains were set to run for 500 million steps and sampled every 1,000 steps (for parameter values) and every 10,000 steps (for trees). We downsampled the set of trees recovered by the MCMC chains to obtain 1,000 trees per chain, sampled at regular intervals between the end of the the burn-in and the end of the entire chain.

Because of the long computation times for the larger datasets (lineages VRC26 and CH103), their MCMC chains were interrupted before 500 million steps had been reached. With the exception of heavy-chain VRC26, interrupted chains had ESSs close to or greater than 200 for most parameters and for the likelihood, prior and posterior probabilities. Although MCMC chains failed to converge for the heavy chain data of lineage VRC26, estimates of the parameters of interest (mean mutability changes and fraction of mutability losses across the tree) were numerically close across replicate MCMC chains, and we therefore present the results in the main text.

To investigate their ability to detect consistent changes in mutability, we repeated the phylogenetic analyses on alignments simulated using different population genetic models (Supplementary Methods). For alignments simulated under a model where all sites are equally likely to mutate, mutability losses were estimated to be approximately as frequent as gains. As expected, mutability losses were estimated to be more frequent than gains in datasets simulated under an S5F-based model, where motifs with high S5F score have a high probability of mutating.

### Simulations of sequence evolution on MCC trees

For each branch on an MCC tree, we compared the nucleotide sequences of the parent and descendant nodes to identify the number of codon sites with non-synonymous and synonymous differences. Starting from the parent sequence, we simulated an alternative descendant sequence constrained to have 1) the same number of non-synonymous and synonymous differences from the ancestor and 2) the same amino acid sequence as the true descendant.

Each simulation was done in two steps. First, for each codon site with anon-synonymous difference, we identified all codons coding for same amino acid and with the same number of nucleotide differences from the ancestral codon as the true descendant codon. We used different models to compute the probability of each possible descendant codon (including the true descendant) replacing the ancestral codon. Let *A* be the ancestral codon and *D* one of the possible descendant codons. For the case where *A* and *D* differ by a single nucleotide, we define the transition rate from *A* to *D* as:

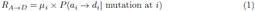

where *i ∈* {1, 2,…, *L*} is the position where codon *A* differs from codon *D*, *L* is the length of the nucleotide sequence, *μ*_*i*_ is the mutation rate at position *i*, and *a*_*i*_ and *d*_*i*_ are the nucleotides at position *i* in *A* and *D*, respectively. For the case where the transition from *A* to *D* requires more than one nucleotide substitution, we define *R*_*A→D*_ as the product of the right-hand side of Equation (1) applied to all intermediary one-nucleotide steps between *A* and *D*, summed across all the possible orders in which the required mutations can be introduced. For example:

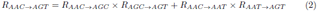

We simulate the evolution of a descendant sequence by sampling a codon from the set of possible descendant codons according to their transition rates from the ancestral codon. The probability of sampling codon *D*_*k*_ is given by:

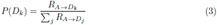

where *k* and *j* index the possible codons.

After simulating evolution at all codon sites where non-synonymous substitutions ocurred, we introduced the same number of synonymous mutations as the number of synonymous codon differences between the ancestral sequence and the true descendant sequence. We allowed synonymous mutations to occur at any codon site except for sites where non-synonymous mutations occurred. To introduce each synonymous mutation, we randomly sampled a nucleotide site, replaced its nucleotide, and either kept the mutation (if it was synonymous) or rejected it (if it was non-synonymous). To simplify implementation, we did not allow for synonymous differences involving more than one nucleotide difference to be simulated – once a codon site was hit by a synonymous mutation, nucleotide sites in that codon could no longer be sampled for subsequent mutations. On average across all datasets, 99.1% of inferred synonymous changes involved a single nucleotide difference (min. 96.1%, max. 99.8%).

Using different models to parameterize Equation (1) and to sample sites for synonymous mutations, we created scenarios where the mutation rate at a site either depends on the nucleotide motif centered at that site, or is independent of nucleotide motifs (see below). To model nucleotide transitions, we made use of the fact that, in addition to estimating the relative mutation rate of different motifs, the S5F model includes the probability of transitions between nucleotides, given the motif where a mutation occurs. All simulations used the S5F-based nucleotide transition probabilities from [21], regardless of whether they assumed variable or constant mutation rates across motifs.

To simulate sequence evolution under an S5F-based model, we let *μ*_*i*_ in Equation (1) be given by:

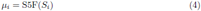

where S5F(*S*_*i*_) is the S5F-score of the five nucleotide motif *S*_*i*_ centered at *i*, from [21].

In addition to the S5F-based model, we performed simulations under two models that assume mutation rates do not depend on the local nucleotide sequence around each site. First, we simulated sequence evolution under a uniform model where all sites are equally likely to mutate (*μ*_*i*_ = 1, regardless of the nucleotide occupying *i* and the neighboring positions). Second, we used a codon-position-based model, where codon positions 1, 2 and 3 have different mutation rates. Estimates of the relative mutation rates of each codon position were obtained from the robust counting inference performed in BEAST and used to parameterize the simulations.

## Acknowledgements

This work was supported in part by the National Institute of Allergy and Infectious Diseases (NIAID) of the National Institutes of Health National (DP2AI117921) and by a Complex Systems Scholar Award from the James S. McDonnell Foundation. The content is solely the responsibility of the authors and does not necessarily represent the official views of the NIAID or the National Institutes of Health. This work was completed in part with resources provided by the University of Chicago Research Computing Center. Code implementing the analyses and figures is available at <u>https://github.com/cobeylab/evolution_of_mutability</u>.

## Supplementary information

### Simulation of B cell receptor alignments

To assess the power of a random local clock model [40] combined with robust counting [39] to detect changes in synonymous and non-synonymous substitution rates during B cell evolution, we used those mehtods to analyze B cell alignments simulated under different scenarios of change in the underlying mutation rates:

**Scenario 1**: uniform mutability, constant mutation rate.

**Scenario 2**: uniform mutability, decreasing mutation rate.

2a: 10% decrease in rate over simulation period

2b: 20% decrease in rate

2c: 30% decrease in rate

2d: 40% decrease in rate

2e: 50% decrease in rate

**Scenario 3**: context-dependency in mutation rates and / or transition probabilities.

3a: S5F context dependency in mutation rates and transition probabilities

3b: hotspot context dependency in mutation rates, uniform transition probabilities, hotspots 3x more likely to mutate

3c: hotspot context dependency, hotspots 30x more likely to mutate For the simulations, we modified a simple forward-time Wright-Fisher model to impose a fitness cost on non-synonymous mutations. The status of a mutation (synonymous or non-synonymous) is defined relative to a fixed reference sequence. As a reference sequence, we used the heavy chain sequence at the root node of lineage CH103, inferred under a logistic growth prior (98-100% identical to sequences inferred in four replicate chains under a constant population size prior). An initial population is generated consisting of a single copy of the reference sequence. At each subsequent generation *t*, *N*_*t*_ sequences are produced by replicating sequences from the previous generation with the possibility of mutation. We assumed *N* grows logistically:

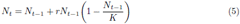

with *N*_0_ = 1 and *N*_*t*_ rounded to the nearest integer. Logistic growth assumes the B cell population initially expands exponentially and then saturates at the carrying capacity *K*.

The probability that a newly generated sequence at generation *t* descends from sequence *i* in generation *t -* 1 is equal to the fitness of sequence *i*, *w*_*i*_, normalized by the sum of fitness values across the entire population at *t -* 1. Any sequence *i* whose amino acid translation is the same as that of the reference sequence has fitness *w*_*i*_ = 1. Each non-synonymous mutation relative to the reference sequence adds *s* to the value of *w*_*i*_, with negative values of *s* representing a fitness cost (*w*_*i*_ cannot go below 0). Newly generated sequences undergo mutations at fixed or variable rates, depending on the scenario.

### Modeling relative and absolute mutation rates

Different models have been proposed to describe variation in somatic hypermutation rates across different nucleotide motifs [7, 21, 22]. For site *i* in sequence *j*, those models can be used to assign a *relative* mutation rate *m*_*i,j*_ to site *i*, based on its local sequence context in sequence *j*. However, the precise relationship between the *relative* mutability of a site and the site’s *absolute* mutation rate is unclear. To model absolute mutation rates as a function of the relative mutability of a sequence’s motifs, we assume that the average mutation rate per site per generation for sequence *j*, 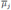, is proportional to the sequence’s average relative mutability, 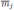

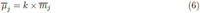

Let 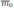 be the average relative mutability of the reference sequence. We choose *k* so that the reference sequence has an average mutation rate per site per generation, 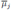 equal to 1*/*4*L*, where *L* is the sequence length (in number of nucleotides):

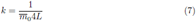

In preliminary simulations under default values for other parameters (see below), this choice of 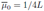 produced alignments visually similar to those observed for real B cell lineages in terms of overall nucleotide diversity. For any sequence *j*, the average mutation rate per site per generation is then given by:

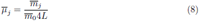

Thus, a sequence with half the average relative mutability of the reference sequence has half the average mutation rate per site per generation. Finally, site-specific mutation rates per generation for sequence *j* are given by:

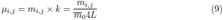

Note that scaling site-specific mutation rates to 1*/*4*L* does not change the relative mutability of the sites. For example, for sites *a* and *b* in the same sequence *j*:

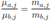

To models scenarios 1 and 2 (where mutation rates are independent of sequence context) we let *μ*_*i,j*_(*t*) = *c*(*t*)× 1*/*4*L* be the mutation rate per time per generation for all sites in all sequences at generation *t*, where 0≤*c*(*t*) ≤1. In scenario 1 we let *c*(*t*) = 1 for all *t*, whereas in scenario 2 we choose decreasing values of *c*(*t*) for different time intervals

In the S5F-based parameterization (scenario 3a), we set *m*_*i,j*_ to the relative mutability scores from [21], based on a five-nucleotide window centered on site *i*. The first two sites and the last two sites, for which S5F is indeterminate since the neighbors are unknown, are assigned mutability zero. We used motif-specific transition probabilities between nucleotides inferred by the S5F model along with motif-specific mutation rates.

In the “hotspot” parameterization (scenarios 3b-c), we let *m*_*i,j*_ be either 1, if site *i* is not at the central position of a hotspot, or *h*, if it is. Site *i* is at the central position of a hotspot if it is occupied by the underlined nucleotide in a *WR<u>C</u>H* or a *D<u>G</u>YW* motif, where *W* = {*A/T*}, *R* = {*A/G*}, *H* = {*A/T/C*}, *D* = {*A/T/G*} and *Y* = {*C/T*} [7]. A site is assigned mutability 1 f it cannot be determined whether or not that site is at the center of a WR<u>C</u>H or a D<u>G</u>YW motif (for example, the left neighbors of the first site in a sequence and the right neighbors of the last site in a sequence are unknown). We assumed uniform transition probabilities between nucleotides.

We ran all simulations with*s* = -0.01, *r* = 0.7 and *K* = 1000 for 2000 generations, sampling 25 sequences every 250 generations starting at generation 500. We decreased *c*(*t*) by 0.2 every 250 generations starting at generation *t* = 1000 for scenario 2a, by 0.4 every 250 generations starting at generation *t* = 1000 for scenario 2b, by 0.5 every 250 generations starting at generation *t* = 750 for scenario 2c, by 0.8 every 250 generations starting at generation *t* = 1000 for scenario 2d, and by 0.1 every 250 generations starting at generation *t* = 1000 for scenario 2d.

We analyzed the simulated alignments using BEAST v.1.8.2 to test if declines in synonymous and non-synonymous substitution rates are correctly detected in scenarios 2 and 3.

### Changes in substitution rates over time

For each tree in the posterior distributions inferred for observed and simulated alignments, we computed the estimated synonymous and non-synonymous substitution rates for each branch by dividing estimated counts of synonymous and non-synonymous substitutions (obtained by robust counting) by the branch’s length measured in time. For each tree in the posterior distributions of simulated alignments, we estimated pairwise linear regression coefficients between branch times (predictor variable, measured at each branch’s parent node) and total, synonymous and non-synonymous substitution rates (response variables).

## Supplementary figures

**Fig S1.**
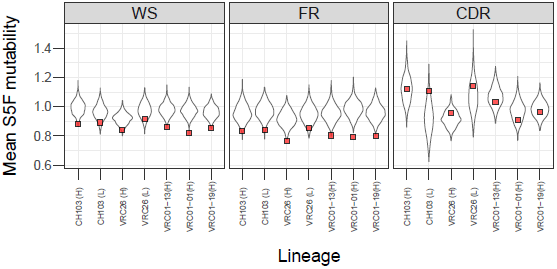
Mutability of the inferred ancestral sequences of long-lived B cell lineages (red squares) compared with the distribution of mutability values obtained by randomizing the ancestral codon sequence while keeping the amino acid sequence constant. Results are shown for the whole sequences (WS) and separately for framework regions (FRs) and complementarity determining regions (CDRs).

**Fig S2.**
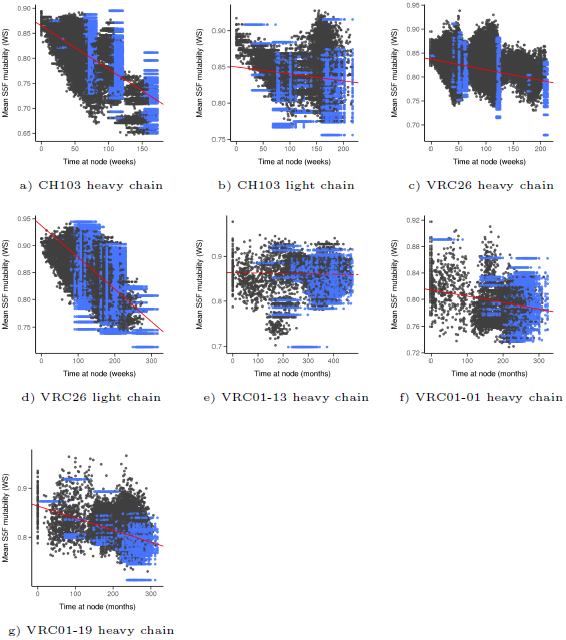
Evolution of S5F-mutability in long-lived B cell lineages. Scatterplots show S5F-mutability over time for nodes from a sample of 100 trees from the posterior distribution. Blue points correspond to terminal nodes (observed sequences), and black points correspond to internal nodes whose sequences were inferred statistically. The red line represents an average of regression lines calculated for each tree in a sample of 1000 trees. Solid lines indicate instances indicate significant relationships (where the highest-posterior density of the slope did not overlap zero), and dashed lines indicate non-significant relationships.

**Fig S3.**
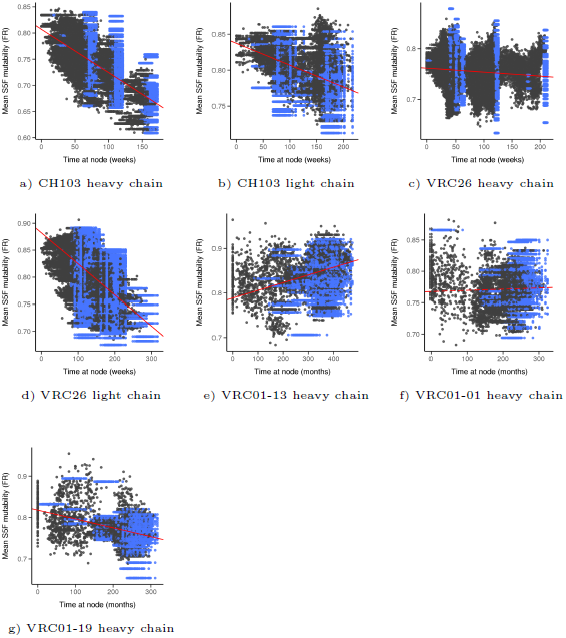
Evolution of S5F-mutability in the framework regions (FRs) of long-lived B cell lineages. Scatterplots show S5F-mutability over time for nodes from a sample of 100 trees from the posterior distribution. Blue points correspond to terminal nodes (observed sequences), and black points correspond to internal nodes whose sequences were inferred statistically. The red line represents an average of regression lines calculated for each tree in a sample of 1000 trees. Solid lines indicate instances indicate significant relationships (where the highest-posterior density of the slope did not overlap zero), and dashed lines indicate non-significant relationships.

**Fig S4.**
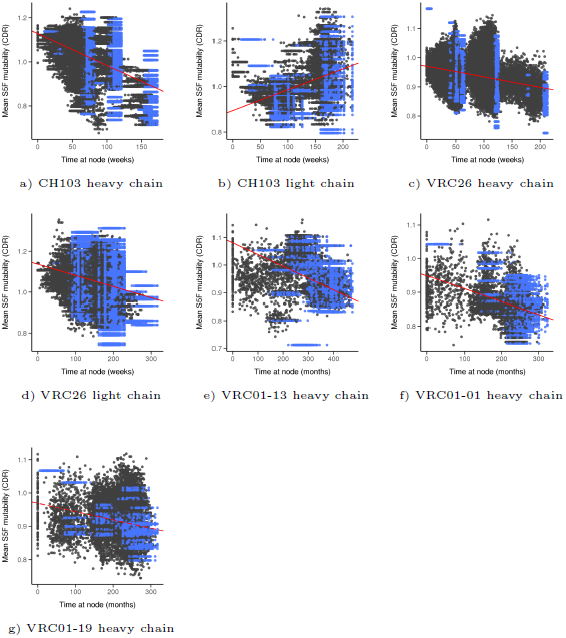
Evolution of S5F-mutability in the complementarity determining regions (CDRs) of long-lived B cell lineages. Scatterplots show S5F-mutability over time for nodes from a sample of 100 trees from the posterior distribution. Blue points correspond to terminal nodes (observed sequences), and black points correspond to internal nodes whose sequences were inferred statistically. The red line represents an average of regression lines calculated for each tree in a sample of 1000 trees. Solid lines indicate instances indicate significant relationships (where the highest-posterior density of the slope did not overlap zero), and dashed lines indicate non-significant relationships.

**Fig S5.**
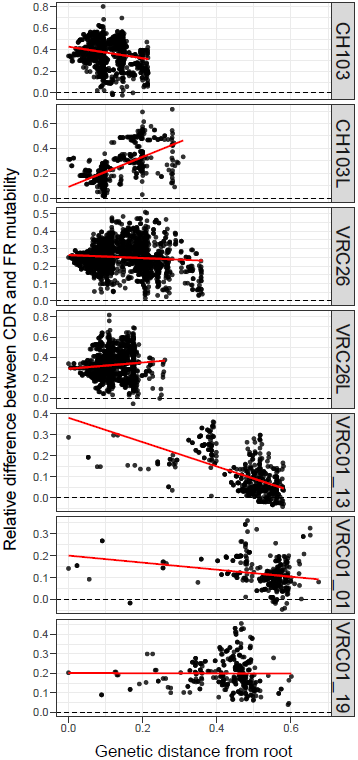
Evolution of the difference in mutability between complementarity determining regions (CDRs) and framework regions (FRs) in long-lived B cell lineages. The relative difference is calculated as the average mutability of CDRs minus the average mutability of FRs, divided by the average mutability of FRs. Each point corresponds to a node in the maximum-clade-credibility tree of each lineage. Linear regression lines are shown in red.

**Fig S6.**
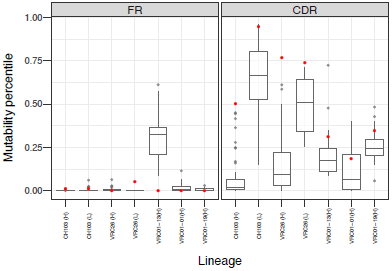
Mutability of B cell receptor sequences from different B cell lineages relative to the expected distribution of mutability values obtained by randomizing codon sequences while keeping the amino sequences constant. The distribution of mutability percentiles obtained for sequences sampled at the last sampling time point in each dataset is shown in gray. The mutability percentile of each lineage’s ancestor is shown in red. Results are shown separately for framework regions (FR) and complementarity determining regions (CDRs).

**Fig S7.**
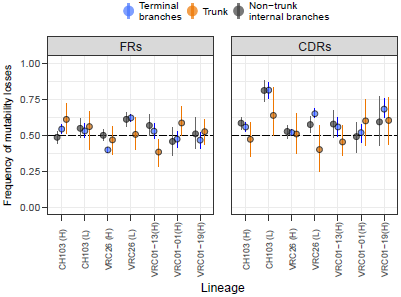
Frequency of mutability losses relative to the total number of changes in S5F mutability caused by synonymous substitutions during the evolution of anti-HIV B cell lineages. Blue indicates changes that on terminal branches, orange indicates changes along the trunk of the tree, and black indicates changes on the remaining internal branches. Results are shown separately for framework regions (FRs) and complementarity determining regions (CDRs). Each point denotes the frequency of changes in mutability that were losses, averaged across a sample of 1000 trees from the posterior distribution. Vertical lines indicate the 95% highest-posterior density interval.

**Fig S8.**
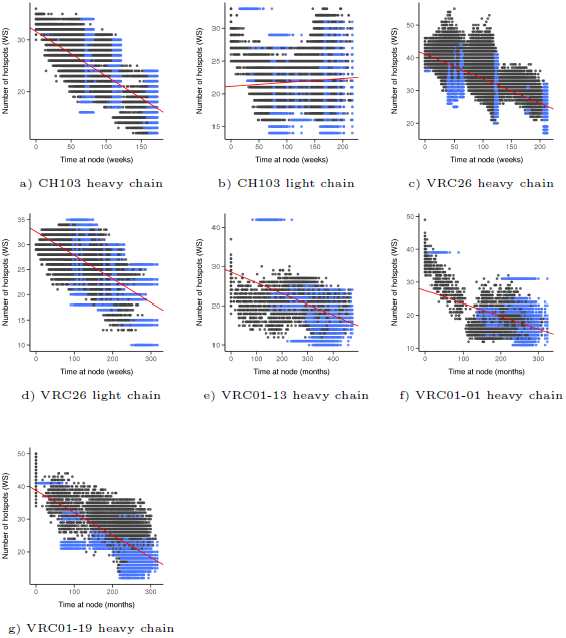
Evolution of the number of WRCH/DGYW hotspots in long-lived B cell lineages. Scatterplots show the number of hotspots over time for nodes from a sample of 100 trees from the posterior distribution. Blue points correspond to terminal nodes (observed sequences), and black points correspond to internal nodes whose sequences were inferred statistically. The red line represents an average of regression lines calculated for each tree in a sample of 1000 trees. Solid lines indicate instances indicate significant relationships (where the highest-posterior density of the slope did not overlap zero), and dashed lines indicate non-significant relationships.

**Fig S9.**
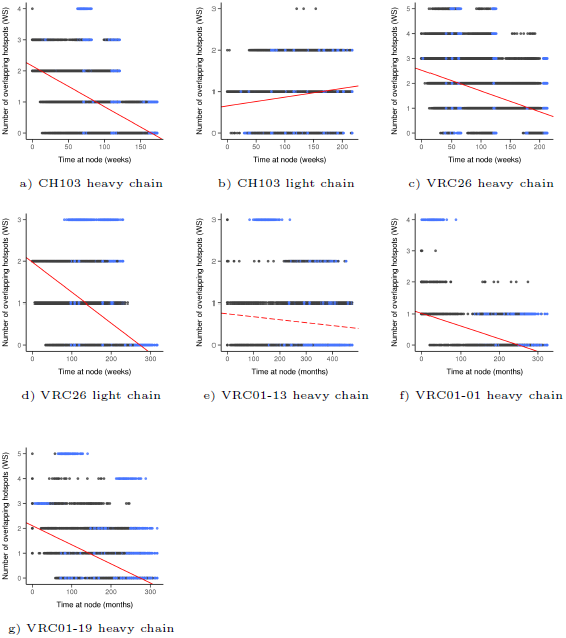
Evolution of the number of overlapping hotspots in long-lived B cell lineages. Scatterplots show the number of overlapping hotspots over time for nodes from a sample of 100 trees from the posterior distribution. Blue points correspond to terminal nodes (observed sequences), and black points correspond to internal nodes whose sequences were inferred statistically. The red line represents an average of regression lines calculated for each tree in a sample of 1000 trees. Solid lines indicate instances indicate significant relationships (where the highest-posterior density of the slope did not overlap zero), and dashed lines indicate non-significant relationships.

**Fig S10.**
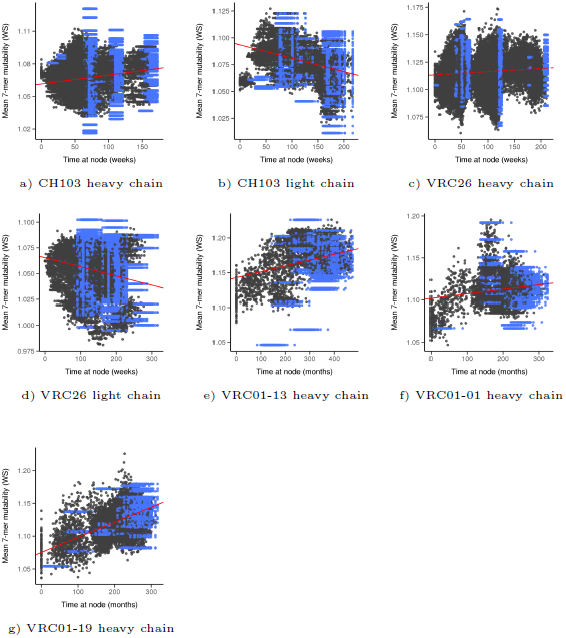
Evolution of 7-mer mutability in long-lived B cell lineages. Scatterplots show 7-mer mutability over time for nodes from a sample of 100 trees from the posterior distribution. Blue points correspond to terminal nodes (observed sequences), and black points correspond to internal nodes whose sequences were inferred statistically. The red line represents an average of regression lines calculated for each tree in a sample of 1000 trees. Solid lines indicate instances indicate significant relationships (where the highest-posterior density of the slope did not overlap zero), and dashed lines indicate non-significant relationships.

**Fig S11.**
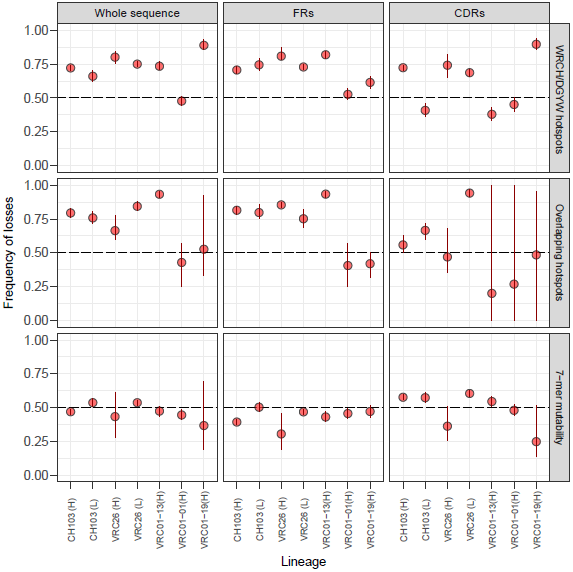
Frequency of mutability losses relative to the total number of changes in mutability during the evolution of anti-HIV B cell lineages. Rows correspond to different mutability metrics, and column contain results obtained for the whole analyzed region of the BCR sequence, and separately for framework regions (FRs) and complementarity determining regions (CDRs). Each point denotes the frequency of changes in mutability that were losses, averaged across a sample of 1000 trees from the posterior distribution. Vertical red lines indicate the 95% highest-posterior density interval.

**Fig S12.**
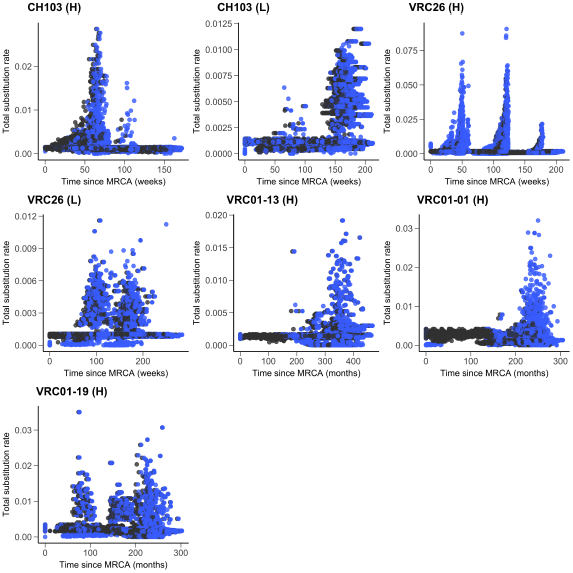
Total substitution rate inferred from the random local clock model, as a function of time for the observed lineages. Each plot shows the points corresponding to a sample of 100 trees from the posterior distribution inferred by BEAST. Terminal branches are shown in blue, and internal branches are shown in black.

**Fig S13.**
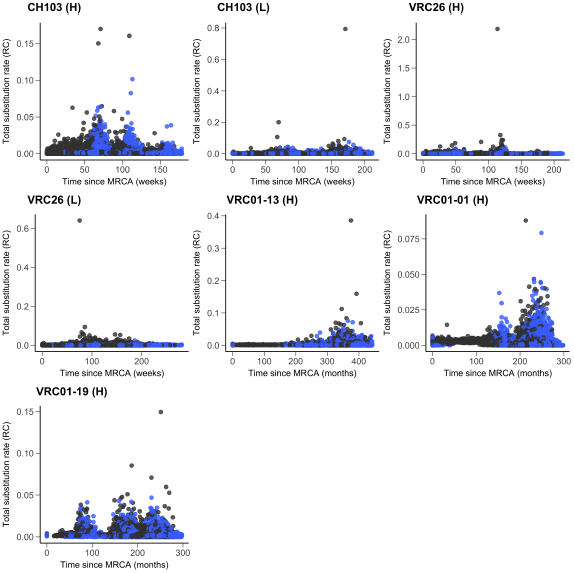
Total substitution rate inferred from robust counting (summing synonymous and non-synonymous rate estimates), as a function of time for the observed lineages. Each plot shows the points corresponding to a sample of 100 trees from the posterior distribution inferred by BEAST. Terminal branches are shown in blue, and internal branches are shown in black.

**Fig S14.**
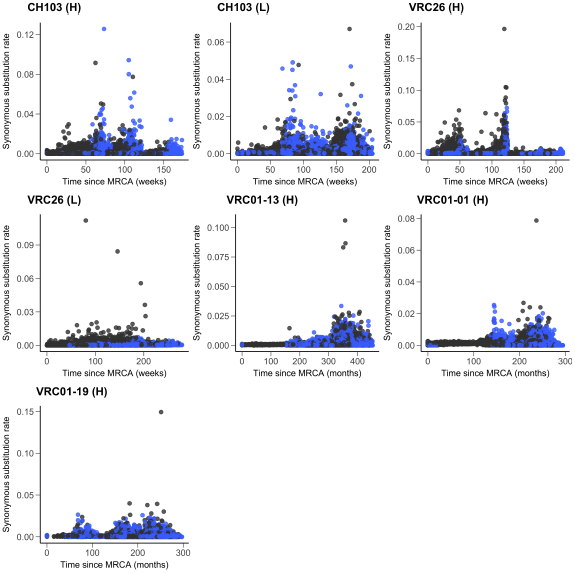
Robust counting synonymous substitution rate as a function of time for the observed lineages. Each plot shows the points corresponding to a sample of 100 trees from the posterior distribution inferred by BEAST. Terminal branches are shown in blue, and internal branches are shown in black.

**Fig S15.**
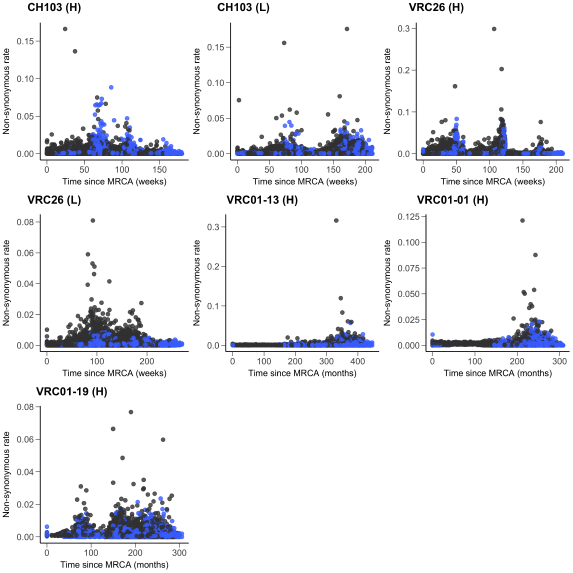
Robust counting non-synonymous substitution rate as a function of time for the observed lineages. Each plot shows the points corresponding to a sample of 100 trees from the posterior distribution inferred by BEAST. Terminal branches are shown in blue, and internal branches are shown in black.

**Fig S16.**
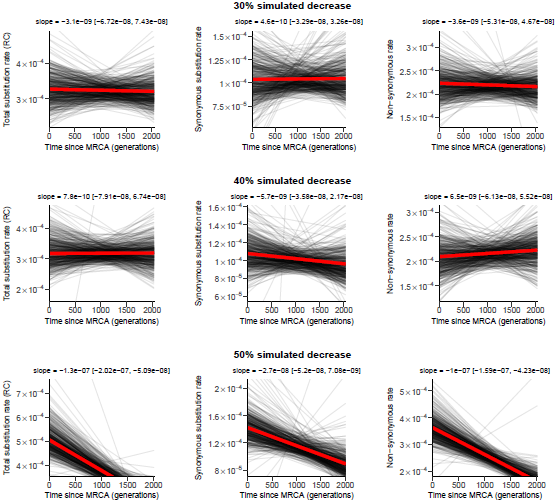
Relationship between robust counting substitution rates and time for simulations performed under different levels of decline in the overall mutation rate. Each black line is the linear regression line between branch-specific rates and times for a single tree from the posterior distribution inferred for a simulated alignment using BEAST. Each plot shows a sample of 500 lines. The red lines are the “average” regression lines, with the average intercept and the average slope calculated from a larger sample of 1000 trees from each distribution.

**Fig S17.**
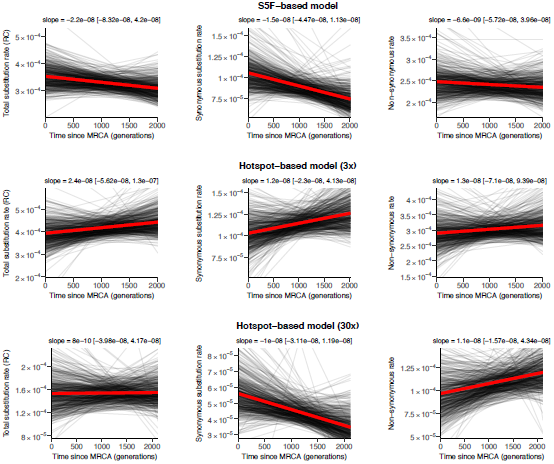
Relationship between robust counting substitution rates and time for simulations performed under models where the mutation rate at each site depends on its S5F mutability or on whether that site is at the center of a WRCH/DGYW hotspot (in which case it mutates either 3 or 30 times more frequently than non-hotspots sites). Each black line is the linear regression line between branch-specific rates and times for a single tree from the posterior distribution inferred for a simulated alignment using BEAST. Each plot shows a sample of 500 lines. The red lines are the “average” regression lines, with the average intercept and the average linear coefficient calculated from a larger sample of 1000 trees from each distribution.

